# Cellular heterogeneity and molecular reprogramming of host response during influenza acute lung injury

**DOI:** 10.1101/2021.08.05.455152

**Authors:** Kai Guo, Dan Justin Kalenda Yombo, Taylor Schmit, Zhihan Wang, Sumit Ghosh, Venkatachelem Sathish, Ramkumar Mathur, Min Wu, Junguk Hur, M. Nadeem Khan

**Affiliations:** Department of Neurology, University of Michigan, Ann Arbor, MI 48109, USA; Department of Biomedical Sciences, School of Medicine and Health Sciences, University of North Dakota, Grand Forks, ND 58202, USA; Abigail Wexner Research Institute at Nationwide Children’s Hospital, Columbus, OH 43215, USA; Department of Pharmaceutical Sciences, School of Pharmacy, College of Health Professions. North Dakota State University, Fargo, ND 58108, USA; Department of Geriatrics, School of Medicine and Health Sciences, University of North Dakota, Grand Forks, ND 58202, USA

**Author notes:** **Dan Yombo and Taylor Schmit contributed equally.**. **Correspondence:** M. Nadeem Khan (Ph.D.), Assistant Professor, Department of Biomedical Sciences, School of Medicine and Health Sciences, University of North Dakota, Kai Guo (Ph.D.), Department of Neurology, University of Michigan, Junguk Hur (Ph.D.), Assistant Professor, Department of Biomedical Sciences, School of Medicine and Health Sciences, University of North Dakota.

## Abstract

Acute lung injury (ALI) caused by influenza A virus (IAV or influenza) manifests from dysregulated cellular interactions between hematopoietic and non-hematopoietic cells that develop into a pathologic host response. However, the lung’s diverse cellular framework that dictates the pathologic host response and acute lung injury remains incompletely understood. We performed a single-cell RNA-seq (scRNA-seq) analysis of total lung cells in mice from severe influenza to examine the cellular heterogeneity and cell-specific regulation of host response. We observed that IAV infection resulted in significant myelopoiesis, predominantly monocyte, and macrophage subsets, which constituted over 50% of total immune cells. IAV infection resulted in the significant loss of endothelial and fibroblast cells, representing the most predominant non-hematopoietic cells and crucial to regulating inflammatory response and barrier integrity. We also show the cell-cell communication dynamics of interferon and chemokine signaling and global regulation of these responses in transition from homeostatic to IAV infection state. These data highlight a robust application of scRNA-seq technology in establishing the atlas of cellular heterogeneity from its homeostatic transition to infection state and the host response regulation in IAV-mediated lung pathology.

## Introduction

Severe influenza disease develops due to exuberant and dysregulated host response that causes acute lung injury (ALI) (1–3). The resolution of influenza requires a coordinated function of leukocyte and non-leukocyte cells that work in tandem to develop an anti-viral host response in the lung. Among leukocytes, resident and infiltrating innate cells mount an early pre-adaptive inflammatory response in the lung. These early immune events are also indispensable to developing antigen-specific T cell responses (4). A significant interdependence and cooperation among the cells of innate and adaptive immune systems are thus required to resolve the infection successfully. However, defects in immune regulation develop a dysregulated inflammatory state that promotes lung pathology and compromises respiratory functions (5, 6).

Despite limitations, mouse models of influenza represent a robust tool to study anti-viral host response and disease pathogenesis, based on several surrogate parameters, such as weight loss, acute lung pathology, and vascular damage (7, 8). Lung resident leukocytes and non-hematopoietic cells initiate a chemokine gradient required to recruit immune cells to the infection site. Although substantial data from mouse and *in vitro* models indicate the significance of leukocyte cells in infection control, how these cells develop a coordinated response that involves either viral clearance or immune-mediated pathology remains to be determined. For instance, while CD8^+^ T cells have long been considered as the most crucial anti-viral immune cells in the lung (9, 10), emerging evidence suggests the pathological role of these cells, which correlate with increased accumulation of CD8^+^ T cells in people with severe flu disease (11, 12). Additionally, CD8^+^ T cells have the potential to cause bystander damage to non-infected cells (13). Similarly, a renewed emphasis on non-epithelial barrier cells, i.e., endothelial and fibroblast cells, highlights the significant role of these cell types in regulating the lung barrier integrity during influenza disease. The feasibility of targeted intervention against influenza requires a comprehensive understanding of the pathologic host response associated with exuberant inflammatory response and acute lung injury. Therefore, a more comprehensive and global understanding of host response to influenza is warranted.

Recent advances in scRNA-seq technology have opened a new way to identify and characterize immune cell landscapes in health and disease (14, 15). The technology can also be used to identify differentially expressed genes and mapping molecular pathways at a single-cell level, which might drive complex pathophysiological responses during various diseases (16). To examine the cellular heterogeneity and regulation of host response at a cellular level, we performed scRNA-seq analysis of cells from mock lungs (homeostatic condition) and influenza-infected lungs of severe disease. Our findings reveal that monocytes and macrophages represent the most predominantly recruited and inter-differentiating immune cell types in the lung. An enhanced myelopoiesis with a loss of endothelial and fibroblast cells correlated with acute lung injury and disruption of barrier integrity in our model. Furthermore, the global response regulation and cell-specific communications among hematopoietic and non-hematopoietic cells establish a unique immunoregulatory role of these communications in influenza disease pathogenesis. An enhanced understanding of these interactions is indispensable to the outcome of influenza disease and could likely result in innovative treatments to combat influenza.

## Results

### Macrophages and monocytes predominate as inter-differentiating immune cells in IAV-infected lungs

To interrogate the immune landscapes and cellular heterogeneity associated with severe influenza disease, we performed scRNA-seq from the lungs of mock and IAV (A/PR/8/34 or PR8)-infected mice (Figure 1A). The severe influenza disease was characterized based on weight loss (~20% weight loss), inflammation (cellular infiltration), vascular damage (BALF albumin), and increased susceptibility for secondary bacterial infection by *Streptococcus pneumoniae* (*Spn*) (Figure 1B). Due to the peak of inflammatory response at day 7 post-infection, this time point was chosen for subsequent investigations. The scRNA-seq data were generated with the droplet-based 10x Chromium platform (10× Genomics). We defined inclusion criteria for cells based on observations from the entire dataset, removed low-quality cells accordingly, then performed dimensionality reduction and unsupervised clustering of the 7,672 recovered cells (mock with 4,082 cells and IAV treatment with 3,590 cells) using Seurat (17). The unsupervised clustering grouped the transcriptomes into 27 distinct clusters, visualized by Uniform Manifold Approximation and Projection (UMAP). Using canonical lineage-defining markers to annotate clusters, we defined seven major cell types (Figure 1C), which were further characterized into 17 cell populations (Supplemental Figure 1), and all the cell types were identified in both mock and IAV infected lungs (Supplemental Figure 1). Predicted cell types were validated using established markers for the cell types (Figure 1D), including endothelial cells and fibroblasts (*Pecam1*, *Flt1*, and *Mfap4*), B cells (*Cd79a*, *Ms4a1*, and *Cd19*), T cells (*Cd3g* and *Cd3e*), natural killer (NK) cells (*Nkg7* and *Gzma*), macrophages and monocytes (*Fcgr1*, *Ly6c2*, and *Mafb*), neutrophils (*S100a8* and *S100a9*), and epithelial cells (*Epcam* and *Tmem212*). We then compared the relative distribution of immune cell compartments between the mock and IAV infection groups.

**Figure 1.**
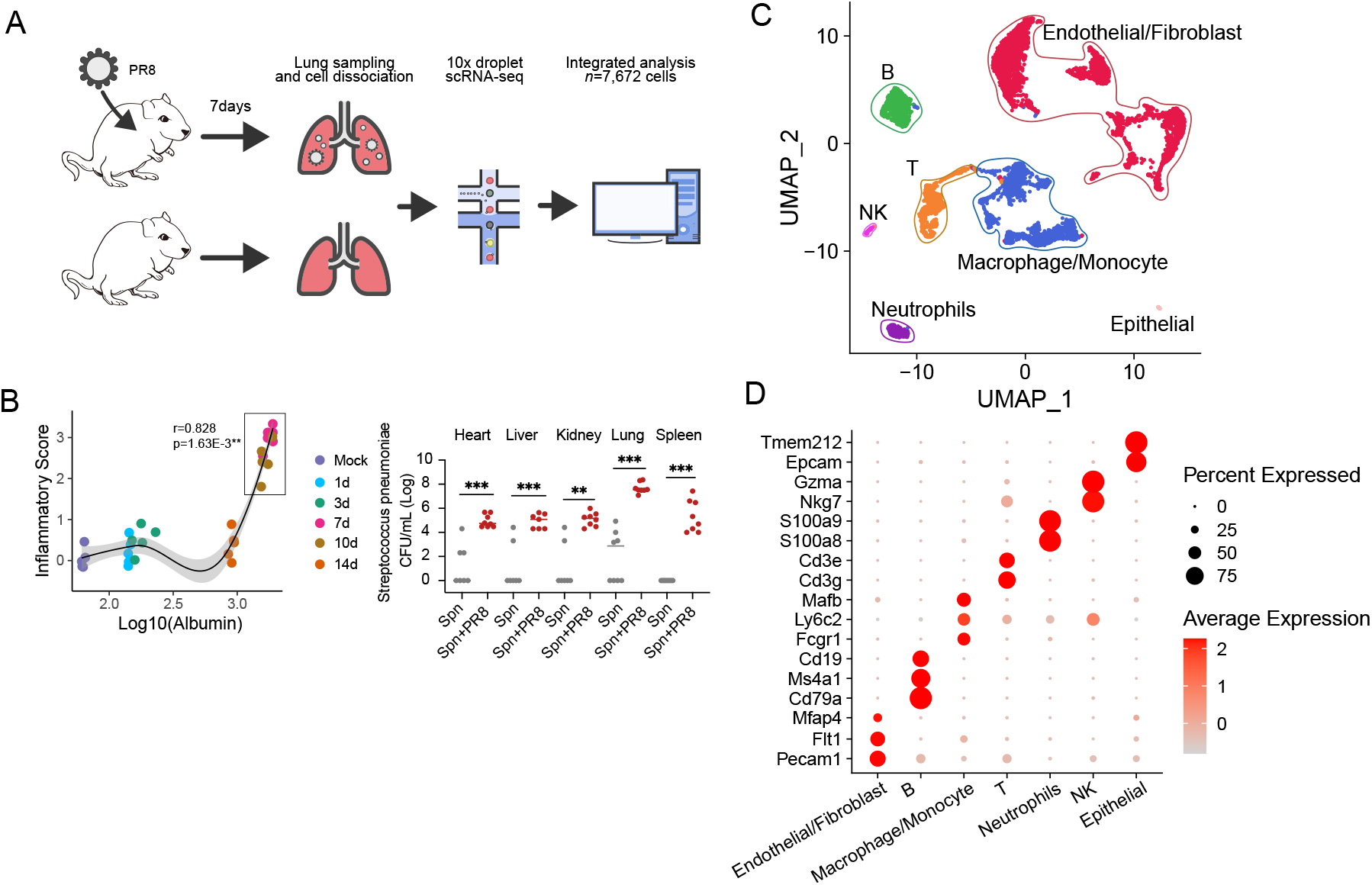
Influenza disease and single cell RNA-Seq landscapes between mock and IAV-infected lungs. Mice were infected with 250 PFU of IAV, BALF and lungs were aseptically isolated for further processing. (A) Schematic representation of experiment flow. (B) Left: Loess regression between log10 (BALF Albumin) and Inflammatory Score (Lung H&E) at 1, 3, 7, 10 and 14 days post-IAV infection, perason correlation were calculated between BALF Albumin and Inflammatory Score at day 7 and day 10; n=5 mice/group, representative of two independent experiments. Right: Bacterial burden in single- (*Spn*) or co-(*Spn* + PR8) infected mice at 48 h post-infection. n=8/group combined of 2 independent experiments, analyzed by two-tailed Student’s *t*-test, **P<0.01, ***P<0.005. (C) Uniform Manifold Approximation and Projection (UMAP) embedding of single-cell transcriptomes from 7,672 cells from mock (n=2) and IAV-infected (n=2) annotated by cell type. (D) Dot plot of mean expression of canonical marker genes for 7 major lineages from tissues of each origin, as indicated.

Myeloid cells in cell clusters belonging to monocytes and macrophages had the most significant increase (Figure 2A, Supplemental Figure 2A). Further analysis revealed that three distinct macrophage cell clusters could be grouped as M1 macrophages, M2 macrophages, and a third macrophage cluster (M0) that could not be identified as M1 or M2 (Figure 2B). Of the three macrophage clusters, only M1 and M2 clusters exhibited the most significant changes upon IAV infection. While IAV infection resulted in a significant increase in M1 macrophage cluster, a simultaneous reduction of the M2 cell cluster was observed (Figure 2A). Besides M1 macrophages, a significant increase in monocyte cluster was observed upon IAV infection (Figure 2A). Together, macrophage and monocyte cell clusters represented over 50% of total leukocytes in IAV-infected lungs. Dendritic cells and neutrophils marginally increased, while the NK cell cluster had no quantitative change (Figure 2A). Therefore, our subsequent analysis and validation focused on macrophages and monocytes due to their predominant response in our IAV model.

**Figure 2.**
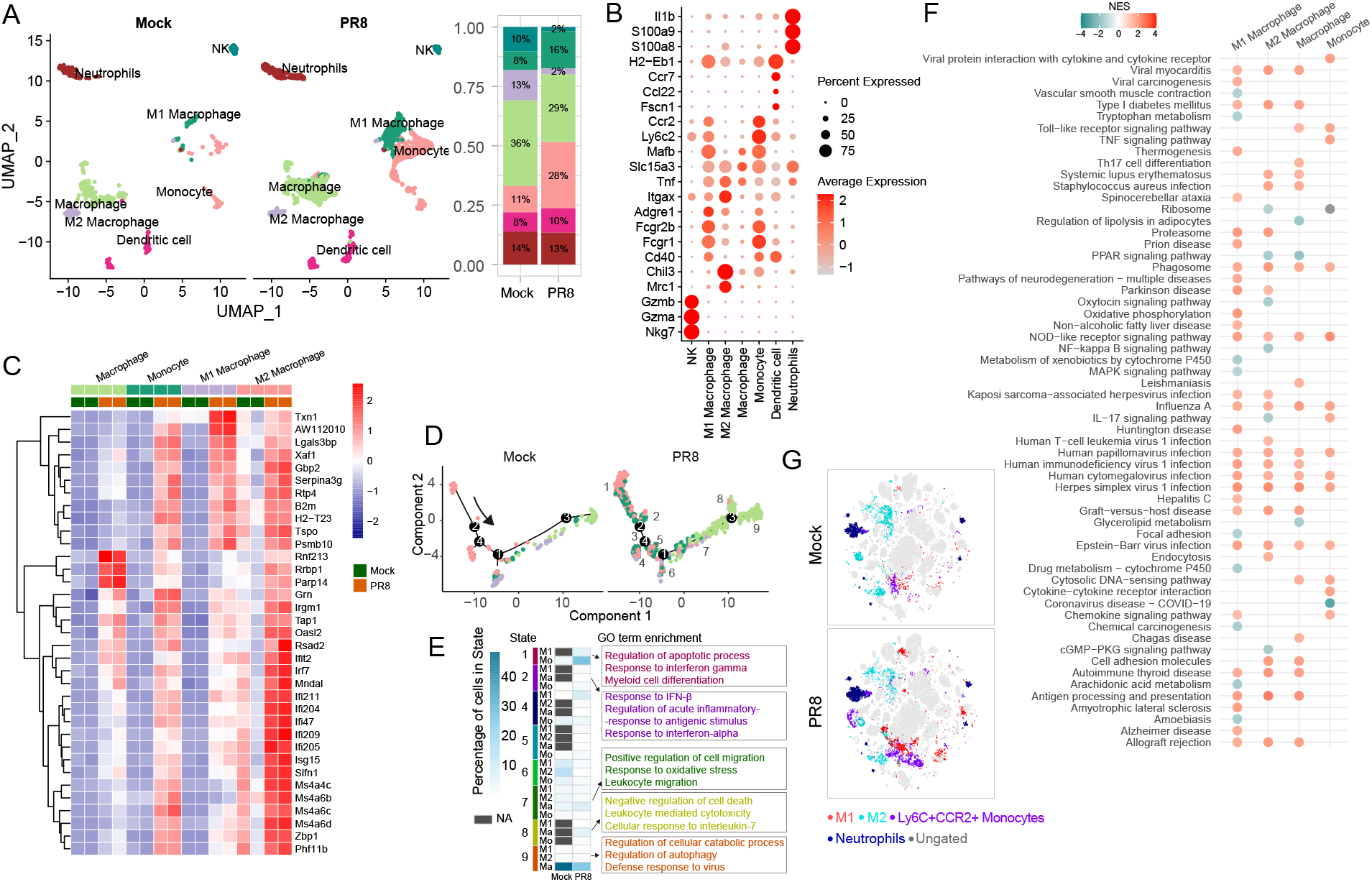
Innate immune cell landscapes in mock and IAV-infected lungs. Mice were infected with 250 PFU IAV, lungs were aseptically isolated and single-cell suspension was prepared for sc-RNAseq and flow cytometry. (A) UMAP plot of innate immune cells, and cell relative proportion of seven subsets (M1/M2 macrophage, macrophage, monocyte, dendritic cells, neutrophils and NK cells) from tissues of each origin, color-coded by clusters and cell subsets as indicated. (B) Dot plot of mean expression of canonical marker genes for 7 cell types, as indicated. (C) Heatmap depicting relative expression of shared differentially expressed genes (DEG) in monocyte, M1 macrophage and M2 macrophage between mock and IAV infection. (D) Pseudotime trajectories of macrophage and monocyte cells in mock and IAV infection along with Seurat cluster. Cells were sorted along the pseudotime. (E) Heatmap comparison of percentage of cells from mock and IAV infection occupying individual states (left) and significantly enriched GO terms from differentially regulated genes of each state (right). Differentially expressed genes were identified as significant at adjusted P-value <0.05 and |avg_logFC| >0.25. Enriched terms were identified as significant at an adjusted P-value < 0.05. (F) Gene Set Enrichment Analysis (GSEA) for KEGG pathways for innate immune cells. Color stands for the up-regulated (red) or down-regulated (blue) after IAV infection. Enriched terms were identified as significant at an adjusted P-value < 0.05. NES: normalized enrichment score. (G) t-SNE diagram of mock and PR8-infected mice (n=5/group representative of two independent experiments) showing M1 (CD11b^+^ CD38^+^ CD163^−^ CD86^+^, iNOS^+^) and M2 (CD11b^+^ CD38^−^ CD163^+^ CD206^+^, Arg-1^+^) macrophages, Inflammatory monocytes (CD11b^+^Ly6C^+^Ly6G^−^CCR2^+^) and neutrophils (CD11b^+^Ly6G^+^) in the lungs at 7 days post-IAV infection.

Flow cytometry-based t-distributed stochastic neighbor embedding (t-SNE) analysis, a dimensionality reduction technique, validated the scRNA-seq findings, demonstrating that monocytes and macrophages were the most significantly increased myeloid cells in IAV-infected lungs (Figure 2G, Supplemental Figure 2B). While the lung IAV-infected mice had significantly increased proportion of M1 and monocytes, M2 and neutrophils showed comparable levels with control mock mice at day 7 post-infection (Supplemental Figure 2B). t-SNE analysis used the conventional markers to establish the phenotypic diversity in monocytes and macrophages: M1 macrophages (CD11b^+^ CD38^+^ CD163^−^ CD86^+^, iNOS^+^), monocytes (CD11b^+^ Ly6C^+^ CCR2^+^), M2 macrophages (CD11b^+^ CD38^−^ CD163^+^ CD206^+^, Arg-1^+^), and neutrophils (CD11b^+^ Ly6G^+^).

Next, we traced cell fate and reconstructed cell lineage direction using the cell trajectory approach to elucidate the relationship between monocyte and macrophage subsets *vis-à-vis* cell differentiation dynamics. Single-cell pseudotime trajectories revealed 9 cell states (no cells in state 3) and 4 branch points (Figure 2D). Parsing the Monocle states by mock and IAV-infected lung cells on the single-cell trajectory revealed that the mock and IAV-infected M1 macrophages occupy multiple overlapping states (Figure 2D). Cells from mock and IAV-infected groups were ordered along pseudotime (Supplemental Figure 2C), and a vast majority of monocytes and M1 macrophages aligned in a continuum along the inflammatory state 1 (Figure 2, D and E), characterized by enrichment of regulation of apoptotic process, response to interferon gamma and myeloid cell differentiation. In contrast, the mock lung in state 1 had fewer monocytes. Note, M1 macrophages arrive only after IAV infection in states 1 and 2. In response to IAV infection, an increased portion of macrophages was also found in states 7, 8, and 9, associated with positive regulation (GO annotations) of cell migration, negative regulation of cell death, and regulation of cellular catabolic process. Strikingly, compared with IAV infected lungs, no macrophages or monocytes were found in state 8 in the lungs of mock mice, which may indicate that state 8 has a unique functional response to IAV infection (Figure 2E). Our data show that upon IAV infection, monocyte and M1 macrophages follow a similar cell fate trajectory based on quantifying the entire transcriptome during the dynamic processes (Figure 2D). Furthermore, monocyte and M1 macrophage exhibited a significant overlap of common differentially expressed genes (DEGs) belonging to host response and inflammation (Figure 2C, Supplemental Figure 2D, Supplemental Table 2). The gene set enrichment analysis (GSEA) for Kyoto Encyclopedia of Genes and Genomics (KEGG) showed that macrophages and monocyte pathways exhibited a significant functional overlap (Supplemental Table 5). Overlaps included specific pathways associated with acute lung injury, oxidative stress, and chemokine signaling, predominantly expressed in the two cell types (Figure 2F). Overall, these data suggest that monocyte and M1 macrophages act as dominant innate cells in IAV-infected lungs and that these cells exhibit a prominent functional and lineage differentiation overlap.

Finally, we assessed the changes in T and B-cell landscapes in the lungs of mock and IAV-infected mice. Similar to innate cells, lineage-defining markers annotated T and B-cells (Supplemental Figure 2E). Our data show that CD4^+^ T cells represented a much larger proportion of lymphoid cells at the homeostatic level. IAV infection induced only a marginal increase in the quantitative frequency of CD4^+^ T cells. IAV infection led to a more robust increase in CD8^+^ T cell frequency (Supplemental Figure 2F). Moreover, the CD8^+^ T cells acquired a potent cytotoxic phenotype 7 days post-IAV infection, characterized by granzyme expression. These data suggest that in primary influenza infection, CD8^+^ T cells evolve into an early effector response that correlates with the resolution of primary infection. In contrast, the delayed CD4^+^ T cell responses might have a crucial role in facilitating antibody responses, protecting against re-infection, or even modulating the CD8^+^ T cell responses against re-infections.

### scRNA-seq identifies IAV-mediated disruption of cellular landscapes of non-hematopoietic lung cells

Increasing evidence suggests that non-hematopoietic cells play a crucial role in regulating pulmonary inflammation and barrier function in infection models (18–21). Using scRNA-seq data, we established the global changes in the atlas of non-hematopoietic cells and their response regulation in the lungs of IAV-infected mice. Lineage-defining markers annotated six distinct cell clusters of non-hematopoietic cells (Figure 3, A and D), i.e., endothelial cells, fibroblasts, myofibroblasts 1, myofibroblasts 2, pericytes, type-II pneumocytes. While all six cell clusters were present in both mock and IAV-infected lungs, endothelial cells represented the most predominant cell type, followed by fibroblasts and pericytes. IAV infection reduced the frequency of all six cellular clusters (Figure 3E). This reduction contrasts with changes observed in immune cells, where IAV infection led to significantly expanding myeloid and CD8^+^ T cells (Figure 2, A and G; Supplemental Figure 2F). To validate scRNA-seq, we performed immunofluorescence staining of lung sections for endothelial (CD31) and fibroblast cells (Col1) (Figure 3, B and C). Consistent with scRNA-seq, immunofluorescence data demonstrate a significant reduction in protein expression of CD31, indicating compromised barrier integrity and reduced endothelial cell presence in IAV-infected lungs (Figure 3C). Consistent with scRNA-seq, we found a significantly reduced number of fibroblasts in the lungs of IAV-infected mice (Figure 3H). However, contrary to flow cytometry and scRNA-seq, immunofluorescence staining of lung fibroblasts revealed an increased expression of collagen-1 (col1) in IAV-infected mice (Figure 3B). Furthermore, immunofluorescent staining of EpCAM^+^ lung epithelial cells also showed damage to the bronchial epithelial layers with loss of continuity in IAV-infected mice (Supplemental Figure 3E). These data suggest that although IAV infection induces a significant loss of fibroblasts, they become functionally more active in producing extracellular matrix proteins. Together, these data show a compromised barrier integrity in IAV-infected lungs; and endothelial cells and fibroblasts representing the most abundant non-hematopoietic cell types in the lung. In addition, endothelial cells and fibroblasts also expressed several shared genes associated with interferon signaling, chemokine signaling, cellular inflammation, and lung barrier regulation (Figure 3F, Supplemental Figure 3, A, B and D, Supplemental Table 4).

**Figure 3.**
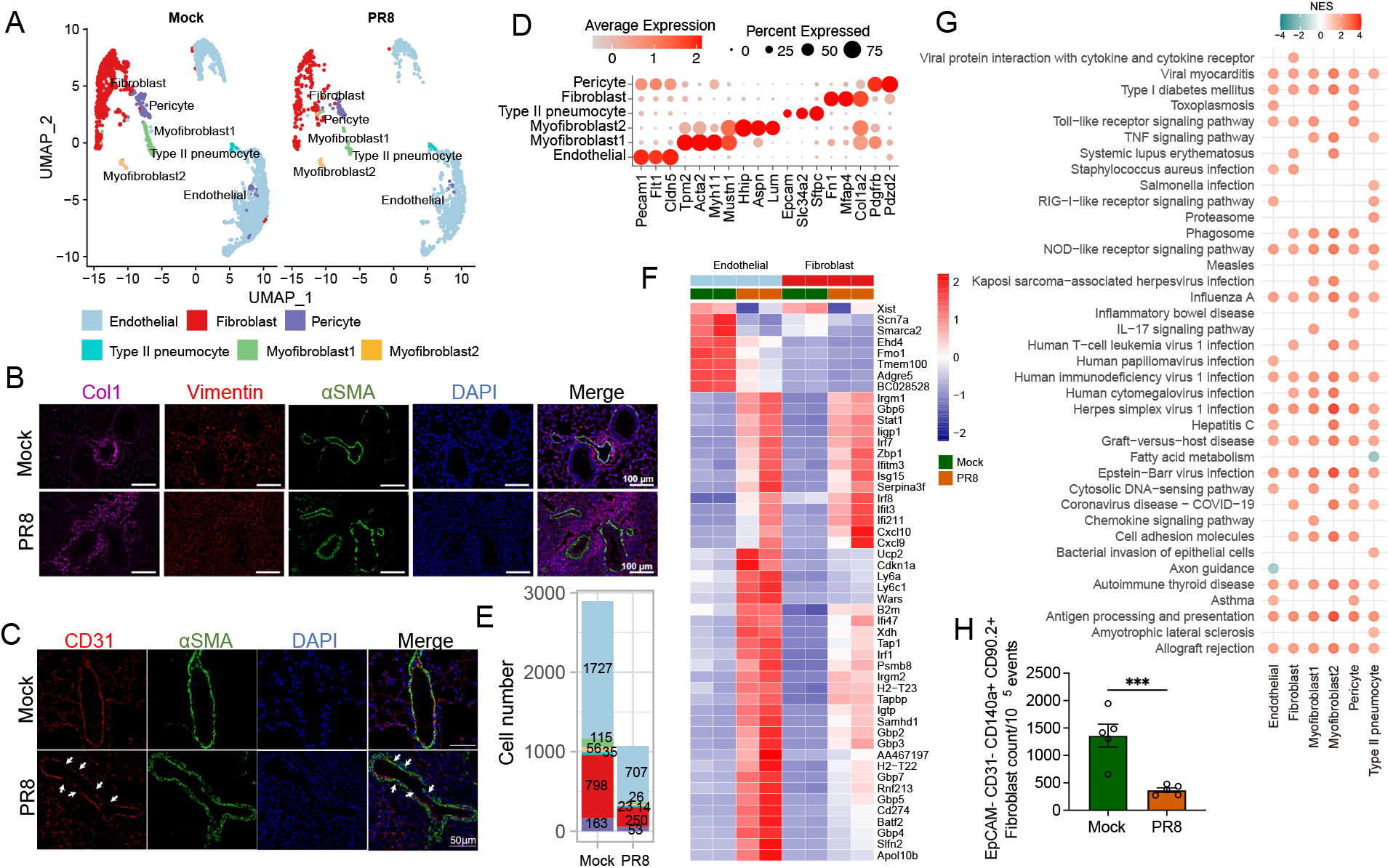
scRNA-Seq landscapes of non-hematopoietic cells in mock and IAV-infected lungs. Mice were infected with 250 PFU IAV. At day 7, mice were euthanized, and lungs were aseptically isolated for immunofluorescent staining or single-cell suspension preparation for sc-RNAseq and flow cytometry. (A) UMAP plot of non-hematopoietic cells color-coded by clusters and cell subsets as indicated. (B&C) Immunofluorescent staining of fibroblasts (B) and endothelial cells (C) in lung sections of mock and IAV-infected mice at day 7 post-infection: (B) Lung fibroblasts were stained with Vimentin(red) and Col1(magenta), airway and vascular smooth muscle cells were highlighted by αSMA(green) staining, and nuclei were counterstained with DAPI (blue). Representative images of 2 different experiments, n=4 per group, images taken at X40 magnification. (C) IAV infection induces a vascular inflammation with damage to the endothelial cells layer integrity, endothelial cells were stained with CD31(Red), smooth muscle cells with αSMA (green), and nuclei were counterstained with DAPI (blue). The loss of vascular endothelial integrity in IAV-infected lung sections, exposing the smooth muscle layer, is highlighted with white arrows. Representative images of 2 different experiments, n=4 per group, images taken at X63 magnification. (D) Dot plot of mean expression of canonical marker genes for 6 cell types, as indicated. (E) Cell numbers of each cell type among mock and IAV-infected lungs. (F) Heatmap depicting relative expression of top 50 shared differentially expressed genes in endothelial and fibroblast between mock and IAV infection. Differentially expressed genes were identified as significant at adjusted P-value <0.05 and |avg_logFC| >0.25. (G) GSEA for KEGG pathways for non-hematopoietic cells. Only top 20 significantly changed canonical pathways in each cell type were shown. Color stands for the up-regulated (red) or down-regulated (blue) after IAV infection. (H) Count of lung fibroblasts/10^5^ events; lung fibroblasts were quantified using FACS staining in lung single-cells prepared from mock and IAV-infected mice at day 7 post-infection. Data shown as means ± SEM, significance was determined by two-tailed Student’s *t*-test. Data is representative of 2 different experiments with n=5, *** P <0.005.

Next, we identified cell-specific differentially regulated pathways in mock and IAV-infected lung cells (Figure 3G) through gene set enrichment analysis (GSEA), which facilitated biological interpretation by robustly detecting concordant differences at the gene set or pathway level (Supplemental Table 6). The top 20 significantly changed canonical pathways in each cell type based on the adjusted P-value are visualized (Figure 3G). Despite a significant functional overlap, a cell-specific molecular regulation pattern manifested among non-hematopoietic cells. For instance, endothelial cells and type-II pneumocytes explicitly downregulated axon guidance and fatty acid metabolism pathways, respectively. The IL-17 signaling was distinctively expressed in myofibroblast following IAV-infection. All cell types commonly upregulated the pathways associated with anti-viral defense (i.e., influenza, HIV-1, herpes simplex virus (HSV)-1 infection, Epstein-Barr virus infection, and NOD-like receptor signaling). These data suggest that while each of the six barrier cells maintains its distinct identity, they share a common identity vis-à-vis regulating inflammation and barrier maintenance. These data also highlight that endothelial cells and fibroblasts represent the most predominant non-hematopoietic cells associated with regulating the lung barrier.

### Molecular reprogramming and cell-cell communications identify the flow of information from homeostatic to IAV infection state

To study the molecular associations underlying cellular communications, we performed cell-cell communication analysis of receptor-ligand pair interactions using CellChat (22). We detected 459 significant ligand-receptor pairs among the 17 cell groups, which were further categorized into 113 signaling pathways, including IFN-I, IFN-II, CD45, IL1, and TGFβ (Supplemental Figure 4, A and B). Comparing the outgoing and incoming interaction strength in 2D space allows for identifying cell populations with significant changes in sending or receiving signals between different datasets. Monocytes and M1 Macrophages emerged from the scatter plot as a significant source and target in IAV-infected lungs compared to mock (Figure 4A).

**Figure 4.**
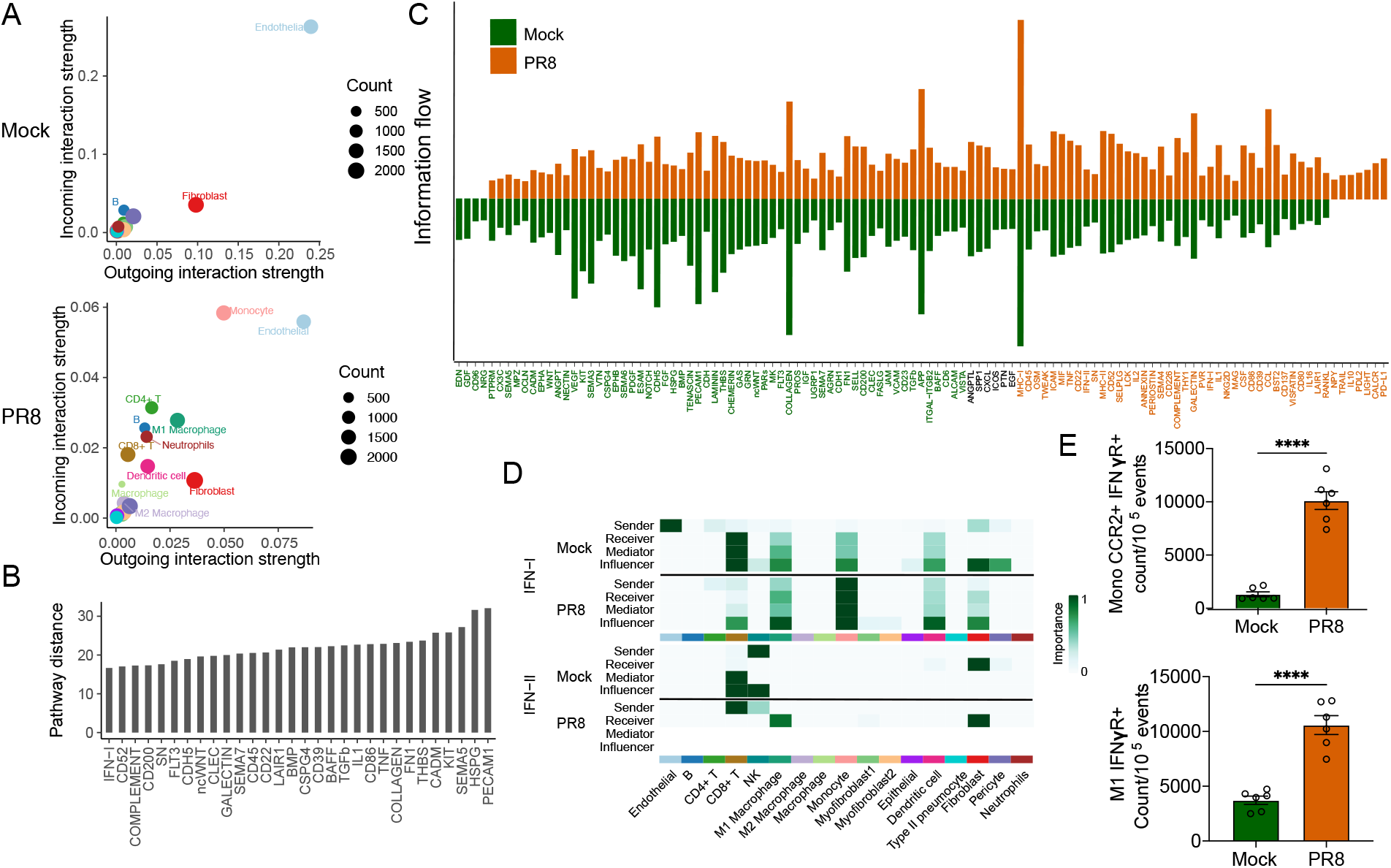
Cell-cell communications between mock and IAV-infected lungs. Mice were infected with 250 PFU IAV. At 7 days post-infection, lungs were aseptically isolated and single-cell suspension was prepared for sc-RNAseq and flow cytometry. (A) Scatter plot show the major source and targets in IAV infection compared to mock. The color indicates the cell types. Dot size is proportional to the number of inferred links (both outgoing and incoming) associated with each cell group. (B) The overlapping signaling pathways between mock and IAV were ranked based on their pairwise Euclidean distance in the shared two-dimensions manifold. Larger distance implies larger difference of the communication network between mock and IAV-infected mice. (C) All the significant signaling pathways were ranked based on their differences of overall information flow within the inferred networks between mock and IAV infection. The signaling pathways colored by red are more enriched in mock, and the ones colored by green were more enriched in IAV-infected mice. (D) Heatmap shows the relative importance of each cell group based on the computed for network centrality measures of IFN-I and IFN-II signaling for mock and IAV infection. (E) The expression of interferon gamma receptor (IFNγR) in CCR2+ monocytes (Top) and M1 macrophages (bottom) was determined in the lung single-cells from mock and IAV-infected mice at day 7 post infection, by FACS; Mono: monocytes (CD11b^+^ Ly6C^+^ Ly6G^−^ cells), M1: M1 macrophages gated as CD11b^+^ CD38^+^ CD86^+^ CD163^−^ iNOS^+^ cells. Data shown as means ± SEM, representive of 2 different experiments. Statistical significance was determined by two-tailed Student’s *t*-test n=5, **** P<0.001.

We then compared cell-cell communication patterns to identify the most dynamically changing pathways between mock and IAV-infected cells. First, we performed joint manifold learning and classified the inferred communication networks based on their topological similarity. We identified four signaling pathway groups (Supplemental Figure 4A), and none of the groups were unique to only one dataset, suggesting that the entire spectrum of communications is represented in both mock and IAV infection data. By computing the Euclidean distance between any pair of the shared signaling pathways in the shared two-dimensions space, we observed a large distance for multiple signaling pathways. The top 30 most dynamically changing pathways between mock and IAV-infected lungs include pathways notably implicated in pattern recognition (GALECTIN, CLEC), inflammation (PECAM, TNF, IL1, CD45, IFN-1, SEMA7, COMPLEMENT), and cellular repair and regeneration (TGFb, COLLAGEN, ncWNT) (Figure 4B). Thus, these pathways exhibit significantly different communication network architectures.

Biological communications, dynamically active during homeostasis and active infection, are central to resolving infection and restoring tissue integrity. To characterize homeostatic versus infection-driven cellular communications, we compared the overall communication probability between mock and IAV infected lung cells (Figure 4C). However, most of these pathways were highly active, albeit at different levels, in mock and IAV-infected lungs, NPY, TRAIL, IL10, PDL2, LIGHT, CALCR, and PD-L1 represented seven core immunoregulatory pathways uniquely activated in IAV-infected lung cells. A significant number of dynamically active pathways transition from mock (dark orange, >50) to IAV-infected state (green, 38) (Figure 4C). Of note, the transition from mock to IAV infection involved developing highly dynamic communications (molecular pathways) implicated in regulating anti-viral response, cell adhesion, inflammation, apoptosis, and tissue repair (Figure 4C). Next, we determined the cell-specific changes in the outgoing signals (levels of ligands) across all significant pathways using pattern recognition analysis (Supplemental Figure 4B). Monocytes and M1 macrophages also exhibited the sources of major differences in outgoing communications between mock and IAV infection (Supplemental Figure 4B), further affirming the significant role of these cells as a mediator of predominant host response in IAV-infected lungs.

Since interferons are crucial to anti-viral defense, we analyzed the global cellular communication dynamics for cell-cell links and effective system-level analyses of the links for IFN-I and IFN-II. Our data demonstrate that while endothelial and CD8^+^ T cells act as significant senders (producers) and receivers for IFN-I signaling in homeostatic conditions (mock), monocytes and M1 macrophages act as the most prominent cells associated with producers, recipients, mediators, and influencers of IFN-I response during IAV infection (Figure 4D). While most IFN-I interactions among mock lung cells were paracrine, the IFN-I interactions among IAV infection lung cells were predominantly autocrine, and to a lesser extent, producing paracrine signaling to fibroblasts, monocytes, dendritic cells, and CD8^+^ T cells. For IFN-II signaling, we found NK cells as a primary cell producing the homeostatic response. At the same time, fibroblasts acted as the only recipient cell for IFN-II response, and CD8^+^ T and NK cells acted as mediators and influencers of IFN-II response at the homeostatic level. However, CD8^+^ T cells were the predominant producers of IFN-II response in IAV infection, while fibroblasts and M1 macrophages were central recipient cells of this response. CD8^+^ T cells were the most predominant cells producing the IFN-II response in IAV infection, and fibroblasts and M1 macrophages were the central recipient cells for the IFN-II response. However, these ligand-receptor interactions only represented the dominant interactions, where both ligand and receptor expression have increased significantly between the homeostatic and IAV infection state.

Since M1 macrophage represented a dominant cell recipient of IFN-II signal and our cell trajectory data demonstrated the monocyte-M1 macrophage differentiation fate, we determined the expression of IFN-II receptor (IFN-γR) on CCR2 monocytes and M1 macrophages between mock and IAV-infected lung cells. Compared to mock lung cells, IAV infection induced a significant increase of IFNγR on CCR2 monocytes and M1 macrophages (Figure 4E). The expression IFNγR by both CCR2 monocytes and M1 macrophages may be the critical component in the overlapping inflammatory functions between inflammatory monocytes and macrophages in IAV-infected lungs.

### Cellular heterogeneity and regulation of chemokine signaling in IAV-infected lungs

Chemokine and chemokine receptor signaling plays an instrumental role in initiating and amplifying host response to resolve the infection. However, the dysregulated chemokine-chemokine receptor axis is also implicated in developing an unrestrained immune response and consequent immune-mediated pathology. Because various hematopoietic and non-hematopoietic cells can produce CCL and CXCL chemokines, we determined the cellular sources of CCL and CXCL chemokine expression, significantly associated with a transition from mock to IAV infection state in the lung. Figure 5A shows the expression levels of genes belonging to four gene families identified as differentially expressed genes (DEGs; adjusted P-value< 0.05 and |avg_log2FC| >0.25 in at least one cell type). Although a number of cells were associated with the production of these chemokines, macrophages and fibroblasts exhibited a significant upregulation of CCL chemokines (*Ccl2*, *Ccl4*, *Ccl5*, and *Ccl7*) upon IAV infection. Macrophages also served as the only significant source of neutrophil recruitment factors, *Cxcl1* and *Cxcl2*, in IAV infected lungs. Several hematopoietic and non-hematopoietic cells, i.e., macrophages, endothelial cells, myofibroblasts, fibroblasts, pericytes, and B-cells, were associated with producing CD8^+^ T-cell recruitment factors, *Cxcl9* and *Cxcl10*, in IAV-infected lungs. Additionally, the receptor genes,*Ccr1*, *Ccr5*, and *Cxcr4* were significantly different between mock and IAV infected lung cells (Figure 5A).

**Figure 5.**
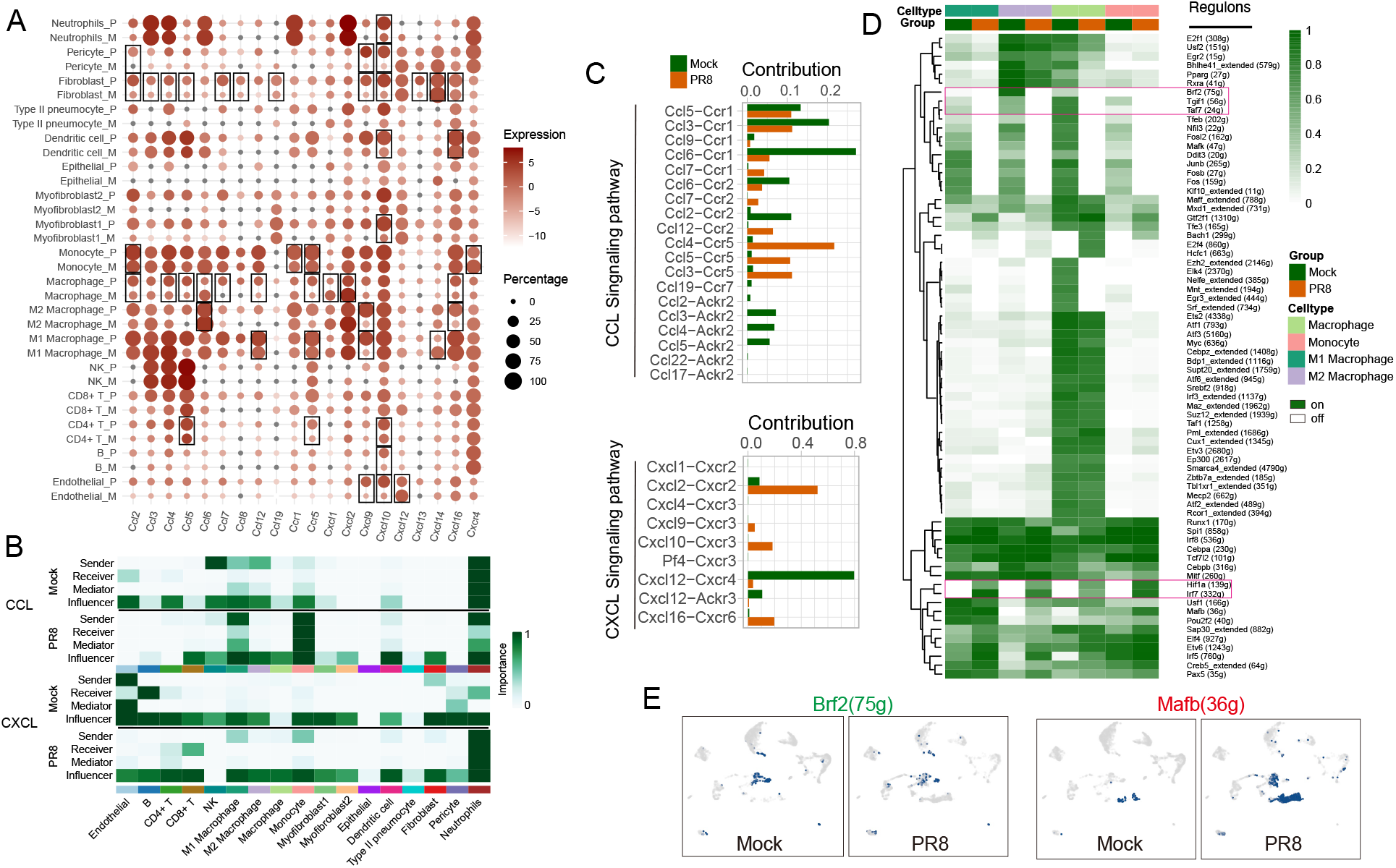
Cellular heterogeneity and host response regulation in mock and IAV-infected lungs. (A) Dotplot show the DEGs from the chemokine and chemokine receptor signaling between in mock and IAV infection. The square indicates that gene was identified as DEGs in specific cell type. (B) Heatmap shows the relative importance of each cell group based on the computed for network centrality measures of CCL and CXCL signaling for mock and IAV infection. (C) Relative contribution of each ligand-receptor pair to the overall communication network of CCL and CXCL signaling pathway among mock and IAV infection. (D) Heatmap of regulon activity analyzed by SCENIC with default thresholds for binarization. The “regulon” refers to regulatory network of TFs and their target genes. “On” indicates active regulons; “Off” indicates inactive regulons. The top rows represent the cell types and the group information. The numbers in parenthesis next to the regulon names indicate the number of genes enriched in regulons. (E) UMAP projection of average binary regulon activity of *Brf2* and *Mafb* in mock and IAV infection.

Next, we investigated the dominant ligand-receptor interactions to the overall communication network of CCL and CXCL signaling pathways between mock and IAV infection regarding their sources and recipient cells for the respective chemokines (Figure 5B). We also investigated the mediators and influencers of the chemokine ligand-receptor communications between mock and IAV infection. Neutrophils and NK cells served as major sources (sender) of the CCL ligands in mock-condition, and IAV infection resulted in a shift from neutrophils/NK cells to monocytes and M1 macrophages. Neutrophils served as the major receiver and mediator of CCL signaling pathway in homeostatic (mock) condition, and monocytes served as a significant receiver and mediator of CCL response in the infection state. The communication networks for CXCL signaling pathways differed substantially from that of CCL signaling pathways. Endothelial cells served as the primary source of CXCL in an autocrine and paracrine manner in mock-infection. In contrast, neutrophils were the most prominent sources, receiver, and mediator demonstrating the significant autocrine signaling in IAV infection. We then focused on the relative contribution of each ligand-receptor to the overall communication network of CCL and CXCL signaling pathways (Figure 5C). Notably, among all known ligand-receptor pairs, the CCL signaling pathway was dominated by the *Ccr1* receptor in mock with *Ccl6*, *Ccl3*, and *Ccl5* ligands. However, *Ccr5* became the primary receptor with *Ccl4*, *Ccl3*, and *Ccl5* as ligands driving the CCL signaling pathway. In the CXCL signaling pathway, the Cxcl2-Cxcr2 ligand-receptor pair was the major contributor to the CXCL communication pathway in IAV infection. In contrast, the Cxcl12-Cxcr4 was the dominant contributor to the CXCL signaling pathway in mock lung cells.

To better understand how IAV infection alters lung global regulatory networks, we implemented the gene regulatory network (GRN) interference approach to investigate the regulons activity of cell-specific transcriptional factors using the single-cell regulatory network inference and clustering (SCENIC) software (23). Based on 300 regulons activity with 10,932 filtered genes with default filter parameters, the selected cells were clustered by cell type, and group information was calculated by SCENIC (Supplemental Figure 5A). Accordingly, regulon activity was binarized and matched with innate cells, and representative regulons and their motifs were listed (Figure 5D). SCENIC revealed that genes regulated by *Hif1a* and *Irf7* were most active in all innate cells following IAV infection (Figure 5D). The *Brf2*, *Tgif1*, *Taf7* genes were slightly reduced from homeostatic to IAV infection-induced transition (Figure 5, D and E). Some transcriptional factors such as *Bach1*, *E2f4*, and *Hcfc1* were primarily associated with macrophage response in IAV infection. Moreover, IAV infection activated significantly more *Stat2* was in endothelial cells after IAV infection, and *Tef* was substantially reduced in IAV infection compared to mock (Supplemental Figure 5, A and B).

## Discussion

Influenza acute pathology manifests from complex global interactions between hematopoietic and non-hematopoietic cells in the lung. These interactions, while crucial to the resolution of infection, can dysregulate host-response and develop a hyper-inflammatory state and concomitant immune-mediated lung pathology. In this transmission, we performed single-cell RNA-Seq from the lung cells of IAV-infected mice that displayed distinct signs of disease and pathology, as demonstrated by a combined metric of lung inflammation, vascular damage, and susceptibility for secondary bacterial diseases caused by *Streptococcus pneumoniae*. Our data indicate that unlike bacterial infections predominated by neutrophil response (24, 25), IAV infection produces a distinct inflammatory response perpetrated by inflammatory monocytes, M1 macrophages, and CD8^+^ T cells (9, 26–29). Endothelial cells and fibroblasts represented the most predominant non-hematopoietic cells, and IAV infection resulted in the significant loss of these cells, impacting the barrier integrity in the lung. We also show the role of IAV in the transition of homeostatic to IAV infection-induced cell-cell communications between hematopoietic and non-hematopoietic cells vis-à-vis interferon signaling in the lung. Finally, our data demonstrate a global regulation of chemokine-chemokine receptor interactions in IAV-infected lungs. Our findings highlight the application of scRNA-seq to establish the atlas of cellular heterogeneity and the regulation of host response in IAV mediated lung pathology.

Unlike bacterial infections that promote CXCL-1/2-mediated neutrophil influx in the lung (30), our data demonstrate that the neutrophils do not act as the predominant cell type infiltrating IAV infected lungs at the peak of the inflammatory response. Instead, monocytes, macrophages, and CD8^+^ T cells dictate the IAV-infected lung environment. Monocytes and CD8^+^ T cells act as anti-viral apparatus (29, 31, 32). Our data importantly demonstrate that monocytes and M1 macrophages are inter-differentiating cells, exhibiting an overlapping effector phenotype. The expression of shared genes and molecular pathways dictating the outcome of inflammatory response in IAV-infected lungs further supports the functional overlap between monocytes and macrophages (Figure 2B & 4E).

The recruitment of monocytes and M1 macrophages also coincided with a significant reduction in the M2 macrophage cell population, further suggesting the suppression of inflammatory restraint caused by M2 macrophages in IAV-infected lungs. These data constitute the first set to denote macrophage heterogeneity and its role in acute lung injury in the IAV model. Our data also proposes cooperativity of inflammatory response between effector CD8^+^ T and myeloid cells, as CD8^+^ T cell effector response correlated with the recruitment of myeloid cells during the peak of acute lung damage in our model (day 7) and diminished CD8^+^ T cell responses associated subsequently with reduced levels of myeloid cells, lung inflammation, and tissue injury. These data deserve further exploration as effector CD8^+^ T cell response can create a bridge for recruiting myeloid cells and modulating their inflammatory function. In our model, we observed only a marginal change in quantitative frequency of CD4^+^ T cells and B-cells, suggesting these responses mature after the peak of the inflammatory response and are potentially crucial to secondary protection against re-infection challenge through antibodies and CD4^+^ T cell-dependent modulation of B-cells and CD8^+^ T cells.

Non-hematopoietic cells in the lung are crucial to the orchestration and regulation of host inflammatory responses during infections and inflammatory diseases (33, 34). However, the relative contribution of each cell type with regard to diverse infections and regulation of inflammatory response is poorly understood. We profiled the non-hematopoietic cells in the lungs of mock-infected lungs and studied the IAV infection-induced quantitative and functional changes to these cells. Among six distinct non-hematopoietic clusters in mock-infected lungs, endothelial cells and fibroblasts represented the most abundant cell types, and IAV infection led to a significant depletion of these cells (Figure 3, A, E, and H). Type-II pneumocytes, pericytes, and myofibroblasts appeared less frequent than endothelial cells and fibroblasts. These data contrast with other organs, such as the gut, where epithelial cells constitute the most abundant cell type (35). The relative abundance of endothelial cells and fibroblasts is crucial to lung function since the lung is a heavily vascularized organ for gas exchange (36, 37), and maintaining vascular integrity is vital to gas exchange and lung function. Endothelial cells and fibroblasts exhibited a spectrum of common DEGs associated with barrier function, and IAV-mediated disruption of barrier integrity was associated with dysregulated interferon signaling. An inflammatory balance that favors the resolution of infection with a concomitant repair is integral to restoring tissue integrity and lung function (38). However, despite much information on influenza host response and inflammation, a knowledge gap remains concerning the dynamically changing pathways during peak tissue inflammation. Our scRNA-seq data show that a number of pathways, involved in pathogen recognition, inflammation, cell-cell adhesion, cellular repair, and regeneration, are most dynamically active between steady-state and IAV infection. Of note, the canonical TGF-β/WNT signaling, involved in cellular proliferation and tissue repair (39, 40), was observed in the top 30 enriched and highly dynamic pathways. Information flow data from cell-cell communication analysis further revealed a downregulation of TGF-β/WNT and Notch signaling during IAV infection state. There is insufficient information on TGF-β/WNT/Notch signaling in influenza ALI, and our data suggest further exploring this axis in flu pathology and tissue repair is warranted.

Ligand-receptor interactions are crucial to cell-cell communication performing the biological function. While upregulation of chemokine/cytokine ligand is a necessary first step in the transmission of biological information, the target recipient cells often exhibit a disparity with the expression of the receptor that receives the biological signal. While some cells might require upregulation of receptor expression to initiate the biological communication, other cells can do that without a necessary change in the expression of the corresponding receptor (41). Our cell-cell communication data show that both type-I and type-II interferons are expressed homeostatically in the lung. However, IAV infection produced a shift in the transition of the source(s) of the cellular response. While some cells acted as recipients of both type-I and type-II interferon responses, monocytes and M1 macrophages significantly enhanced the expression of the receptors for type-I and type-II interferons, demonstrating a dominant ligand-receptor interaction. However, due to the functional overlap between monocytes and M1 macrophages, including the expression of CCR2 on M1 macrophages, we also detected the IAV-infection induced increase in the expression of type-II interferon receptor on monocytes. Nevertheless, whether these ligand-receptor interactions also correspond to producing the biological signals of similar strengths compared to other cell types that did not exhibit a change in receptor expression remains to be determined.

Chemokine gradient formation is essential for coordinated immune cell migration to infected tissues determining the fate of infection resolution (42). However, unrestrained chemokine gradient can also result in dysregulated immune-mediated pathology (43). Our data demonstrate cellular heterogeneity with regard to the expression of CCL and CXCL chemokines, with both hematopoietic and non-hematopoietic cells acting as significant sources of these chemokines in IAV-infected lungs. Our data also suggest that besides senders (ligand producing cells) and receivers (cells expressing corresponding receptors), a number of cells act as mediators and influencers to modulate the intensity of biological communications, regulating the complex biological functions mediated by chemokines and interferons. Since macrophages and monocytes were the most predominant cell types recruited in our model, we studied the changes in cell-specific transcriptional factors and global regulatory networks induced by IAV infection. Our data show a unique pattern of cell-specific regulons with significant overlaps in pathways associated with interferon, HIF-1, and oxidative stress from steady-state to IAV infection state.

Collectively, our data show that scRNA-seq is a powerful tool to study the cellular landscapes and their response regulation during IAV infection. These data demonstrate that IAV infection preferentially promotes the recruitment of myeloid cells while simultaneously depleting non-hematopoietic cells, which are crucial in regulating lung inflammation and barrier integrity. In myeloid cells, we further demonstrated the divergences between IAV and bacterial infection –while neutrophils are perceived as the dominantly recruited cell type in lung bacterial infections, monocytes and macrophages served as the most predominant cell types recruited to IAV-infected lungs. These data led us to postulate that a defect in neutrophil recruitment and the disruption of the lung barrier might explain the heightened susceptibility of IAV-infected lungs for secondary bacterial infections. Finally, our data identified several new cell-specific pathways and regulatory networks that can create a foundation for further exploring the mechanistic relationship of influenza exuberant inflammation with acute lung injury.

## Methods

### Influenza infection model

The C57BL/6 mice were purchased from The Jackson Laboratory and bred in-house. An equal proportion of 6–8-week-old male and female mice were included in all experiments. To establish influenza disease, mice were lightly anesthetized with 4%v/v isoflurane/oxygen mixture and intranasally inoculated with 250 PFU of Influenza A Virus (A/PR/8/34 or IAV or PR8) (Charles River) in 50 μl volume and monitored for signs of morbidity. At indicated timepoints (days 1-14), mice were euthanized by CO_2_ exposure followed by cervical dislocation and processed for downstream applications.

### Histopathology

Lung sections were prepared, processed and stained by the Histology Core, Department of Biomedical Sciences, University of North Dakota. Whole lungs were perfused and fixed with 10% formalin for 24 h before transferring into 70% ethanol prior to processing. Tissues were embedded in paraffin and sectioned into 5 μm sections. Each lung specimen was stained with hematoxylin-eosin and inflammation was evaluated by three pathologists, in a blinded fashion, on a scale of 0–4 with increments of 0.5. For scoring, mock lungs were defined as 0 and lungs with severe cell infiltration and tissue pathology were defined as 4 (44).

### Albumin ELISA

Bronchiolar lavage fluid (BALF) supernatant was preserved at −20 °C until further processing. Albumin concentration in BAL samples was determined using a Mouse Albumin ELISA (ALPCO, Salem, NH) at a 1:10,000 dilution. All samples were read on Synergy HT spectrophotometer, and data were analyzed using KC4 Data Analysis Software (BioTek Instruments).

### IAV-Spn co-infection

*Streptococcus pneumoniae* serotype 6A strain was (BG7322) was obtained from Rochester General Hospital Research Institute and used by others (45). To establish co-infection, mice were intranasally inoccuated with PR8 at day 0 and 2000 CFU of *Spn* 6A in 50 μl sterile PBS at day 7. After 48 h, specimens were homogenized, serially diluted, and plated on blood agar plates. The bacterial CFUs were enumerated next day.

### Flow cytometry

The lungs were aseptically collected from mock and IAV-infected mice and single cells were prepared as previously described (44). One million cells were stained with Ghost Dye-BV510 (Tonbo Biosciences, San Diego, CA) and anti-mouse CD16/CD32 (BD Biosciences, San Jose, CA) at 4°C for 30 min, then stained for surface markers with CD11b-APC/Cy7, Ly6C-BV711, Ly6G-FITC, CCR2-PE/Cy7, F4/80-APC, CD3-APC/Cy7, IFN-γR-BV421 (BD Biosciences, San Jose, CA), CD11c-PerCP/Cy5.5, CD38-FITC, CD86-PE/Cy5, CD206-PE/Cy7, and CD163-BV421, CD8-PE/Cy7, and CD4-BV711 for 30min at 4°C for 30 min. For M1 and M2 macrophages, cells were further fixed and permeabilized using BD CytoFix/cytoperm Kit following the manufacturer’s instructions (BD Biosciences) and stained with iNOs-PE and Arginase1-APC (R&D Systems, Minneapolis, MN). Unless otherwise specified, all fluorochrome-labeled antibodies were purchased from BioLegend (San Diego, CA). To quantify lung fibroblasts, lung single cells were stained with CD45-BV605, CD31-PE, EpCAM-APC, CD90.2-PB, and CD140a-PE/Cy7 antibodies. Lung fibroblasts were gated as CD45^−^ CD31^−^ EpCAM^−^ CD90.2^+^ CD140a^+^ cells as previously described (46, 47). The complete list of antibodies used for flow cytometry (Supplemental Table 1) and the gating strategy for M1, M2 macrophages and fibroblasts can be found in supplemental information (Supplemental Figure 6 & 7).

A BD FACSymphony cytometer was used to acquire 100,000 events and data were analyzed using FlowJo (Tree Star). For the generation of flow cytometry t-Distributed Stochastic Neighbor Embedding (t-SNE) using FlowJO, cells were gated for singlets and live cells before down-sampling to 15,000 events per sample. t-SNE was generated using an iteration number of 1,000, a trade-off of 0.5, and perplexity of 30 (48).

### Immunofluorescence staining

Formalin-fixed and paraffin-embedded lung sections from mock and IAV-infected mice were prepared as previously described (44) and probed for epithelial or endothelial cells detection using anti-mouse αSMA (1:10,000; Sigma-Aldrich, Darmstadt, Germany), rabbit anti-mouse CD31 (1:50; Abcam, Cambridge, UK), or rabbit anti-mouse EpCAM (1:50, Abcam, Cambridge, UK) antibodies. To detect lung fibroblasts, lung sections were stained with rabbit anti-mouse Collagen type I (Col1)( 1:100, AB765P, Sigma Millipore) and chicken anti-mouse Vimentin (1:500, AB5733, Sigma Millipore). Tissues were incubated with corresponding secondary antibodies goat anti-mouse IgG2a-AF488 (1:200; A-21131, ThermoFisher), anti-rabbit IgG-AF546 (1:200; Invitrogen, Carlsbad, CA) or anti IgG-AF633 (A-21070; ThermoFisher), and anti-chicken IgG-AF594 (1:200, A-11042, ThermoFisher) in 5% goat serum/PBS for 1h at RT, nuclei were counterstained with DAPI. All the images were acquired using an Olympus Total Internal Reflection Fluorescence microscope containing a Hamamatsu ORCA-Flash4.0 camera.

### Single-cell RNA-Seq

From single-cell suspensions of mock and IAV-infected lungs, 2×10^6^ cells were resuspended in media containing 10% DMSO and 20% FBS and allowed to slow freeze in Mr Frosty. 3’ single cell gene expression libraries (v3.0) were constructed using the 10× Genomics Chromium system. Single cell library preparation was done by Singulomics Corporation (https://singulomics.com/). A pair-end 150 bp sequencing was performed to produce high-quantity data on an Illumina HiSeq platform(Illumina, San Diego, California, USA).

### scRNA-seq data alignment and sample aggregating

Reads with sequence quality less than 30 were filtered out and then mapped to the GRCm38 mouse reference genome using the CellRanger toolkit (version 3.0.2). Individual sample output files from CellRanger Count were read into Seurat v3 (17) to generate a unique molecular identifier (UMI) count matrix used to create a Seurat object containing a count matrix and analysis. Three quality measures on the raw matrix for each cell: gene number between 200 and 7,000, UMI count above 1,000, and mitochondrial gene percentage below 0.2. Then all data with mock and IAV treatment were integrated for the following analysis. LogNormalize method in Seurat was used for normalizing filtered gene-barcode matrix.

### Dimension reduction, clustering, and visualization

Principal component analysis (PCA) was done by using the top 2,000 most variable genes. To determine the optimal principal component (PC) number, the PC number where the percent change in variation between the consecutive PCs with less than 0.1% was selected. Then Uniform Manifold Approximation and Projection (UMAP) was performed on the principal components for visualizing the cells, and graph-based clustering was performed on the PCA-reduced data for clustering analysis (resolution=0.8) with Seurat v3 (17).

### Cell type annotation and differential expression analysis

To identify the cell type of each cluster, the gene markers for each cluster were identified by using the FindAllMarkers function implemented in Seurat (parameters: min.pct = 0.25, logfc.threshold=0.25). Cell types were assigned by using the CellMarker and PanglaoDB database as the reference, with manual correction. To identify the differentially expressed genes for each type between mock and IAV infected, MAST (49) was used to perform differential analysis. Genes were considered differentially expressed if the adjusted P-value was lower than 0.05 and the average log2FC was greater than 0.25. Kyoto Encyclopedia of Genes and Genomes (KEGG) and Gene Ontology functional enrichment were performed by using richR (https://github.com/hurlab/richR/) package and the adjusted P-value< 0.05 was chosen as the cutoff value to select significant KEGG or GO terms.

### Trajectory inference

The R package Monocle (50) was used to estimate the potential lineage trajectory. Cell trajectory provides a basis for quantitative modeling of cell fates based on gene expression on a time scale (50). The gene-cell matrix in the scale of UMI counts was provided as input to Monocle, and then, its new Cell Data Set function (lower Detection Limit = 0.1, expression Family = nonbinomial.size) was called to create an object. The size factors and dispersion of the gene expression were estimated. We used these subpopulations from the Seurat data set as described previously to identify the DEGs. We then selected the top 1,000 DEGs based on a minimum adjusted P-value of 0.01 for Monocle to order the cells using the “DDRTree” method and reverse graph embedding. Differentially expressed genes for each Monocle state were identified using Seurat.

### Cell-cell communication

To understand global communications among cells, CellChat (22), a tool that can quantitively infer and analyze intercellular communication networks from scRNA-seq data, was used for the cell-cell communication analysis. CellChat predicts major signaling inputs and outputs for cells and how those cells and signals coordinate for functions using network analysis and pattern recognition approaches. First, the significant ligand-receptor pairs among 17 cell groups were identified and categorized into signaling pathways. Second, the key incoming and outgoing signals were predicted for specific cell groups as well as global communication patterns by leveraging pattern recognition approaches. Then, the significant signaling pathways were grouped by defining similarity measures and performing manifold learning from topological perspectives. And the overall communication probability analysis was performed across the mock and IAV-infected datasets.

### Regulatory network inference

The Single-Cell regulatory Network Inference and Clustering (SCENIC) (23) was performed to assess the regulatory network analysis in regard to transcriptional factors and discover regulons in individual cells. Following the standard pipeline, the gene expression matrix with gene names in rows and cells in columns was input to SCENIC (23). The raw count data were extracted from the Seurat object, filtered with the default parameters, finally resulting in 10,932 mouse genes mapped in RcisTarget database (23). The co-expressed genes network for each TF was constructed with GENIE3 software (51), followed by Spearman’s correlation between the TF and the potential targets. Regulons, groups of target genes regulated by a common transcription factor, were generated from the “runSCENIC” procedure based on the correlation between the transcriptional factors and the potential targets. Finally, the regulon activity was analyzed by AUCell (Area Under the Curve) software (23) with the default threshold applied to binarize the specific regulons, and transcriptional factors expressions were projected onto UMAPs.

### Data and software availability

The data were deposited in the NCBI’s BioProject database (PRJNA733762).

### Statistics

Two-tailed Student’s t test was used to asses 2 independent groups with a P value less than 0.05 considered significant. Genes and pathways were considered significant when adjusted P-values (controlled for multiple comparisons using the Benjamini–Hochberg FDR procedure) were <0.05.

### Study approval

Animal care and experimental protocols were in accordance with NIH “Guide for the Care and Use of the Laboratory Animals” and approved by the Institutional Animal Care and Use Committee (IACUC) at the University of North Dakota (Protocol Number 1808-8).

## Author contributions

N.K., J.H. and K.G. designed and supervised the study. K.G. performed single cell data analysis. D.J. and T.S. coordinated sample collection and data interpretation. D.J and T.S. performed flow cytometry, Immunofluorescence staining and all other experiments. K.G., N.K., D.J., T.S., Z.W., and J.H. contributed to the data interpretation. K.G., N.K., D.J., T.S., and N.K. wrote the manuscript, and all authors contributed to the writing and provided comments.

## Acknowledgments

This work was supported by the National Institutes of Health Grant, R01 AI143741 to M. Nadeem Khan. We thank Melody Davis, Abigail Wexner Research Institute at Nationwide Childrens Hospital, Columbus,OH, for critical reading of the manuscript. Figure 1A was created by modifying illustrations provided by Vecteezy.com

---

**Supplemental Figure 1.**
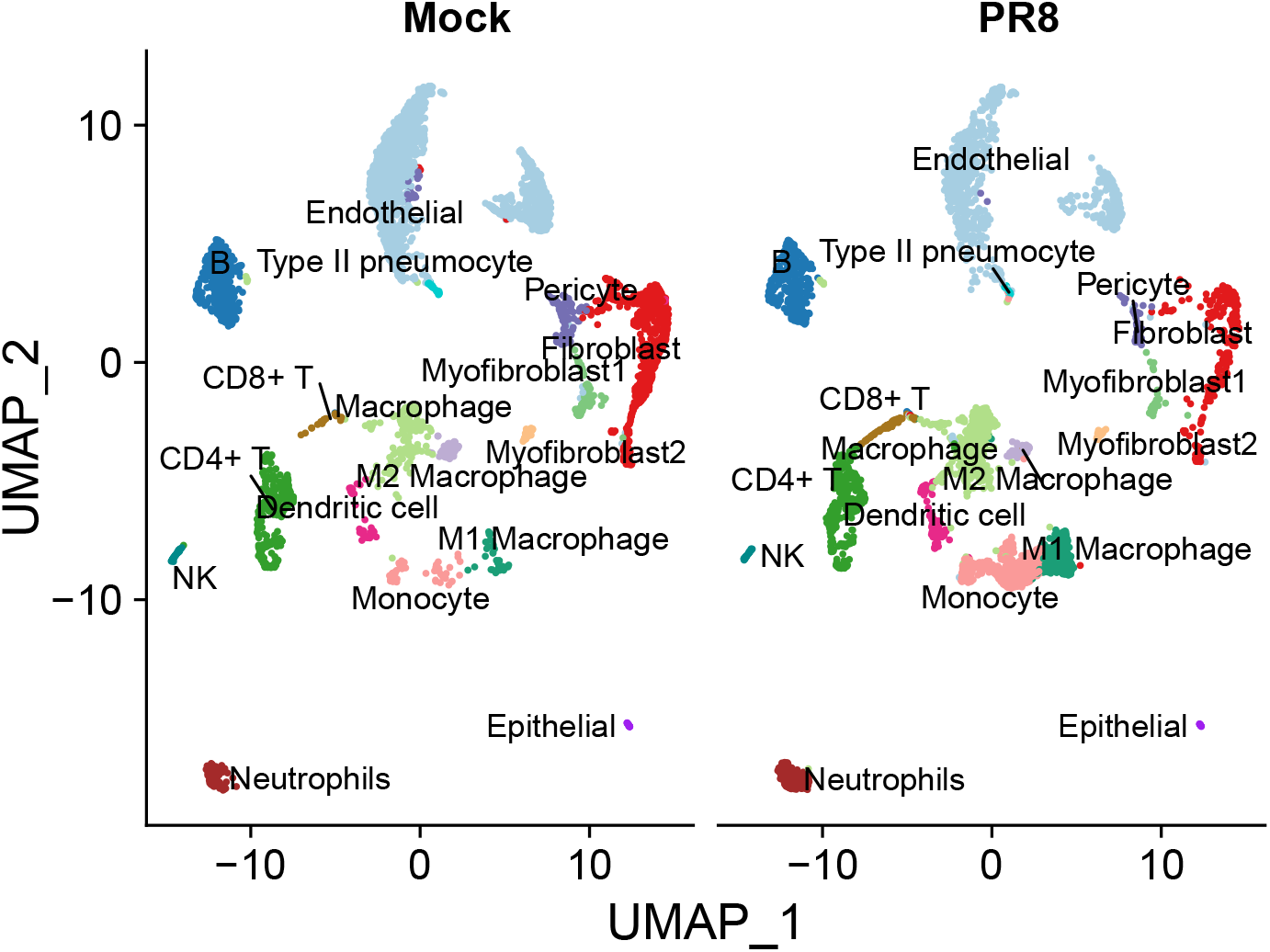
UMAP projection embedding of single-cell transcriptome from mock and IAV-infected lungs. Mice were infected with 250 PFU IAV. At 7 days post-infection, lungs were aseptically isolated and single-cell suspension was prepared for sc-RNAseq. Color stands for cell types.

**Supplemental Figure 2.**
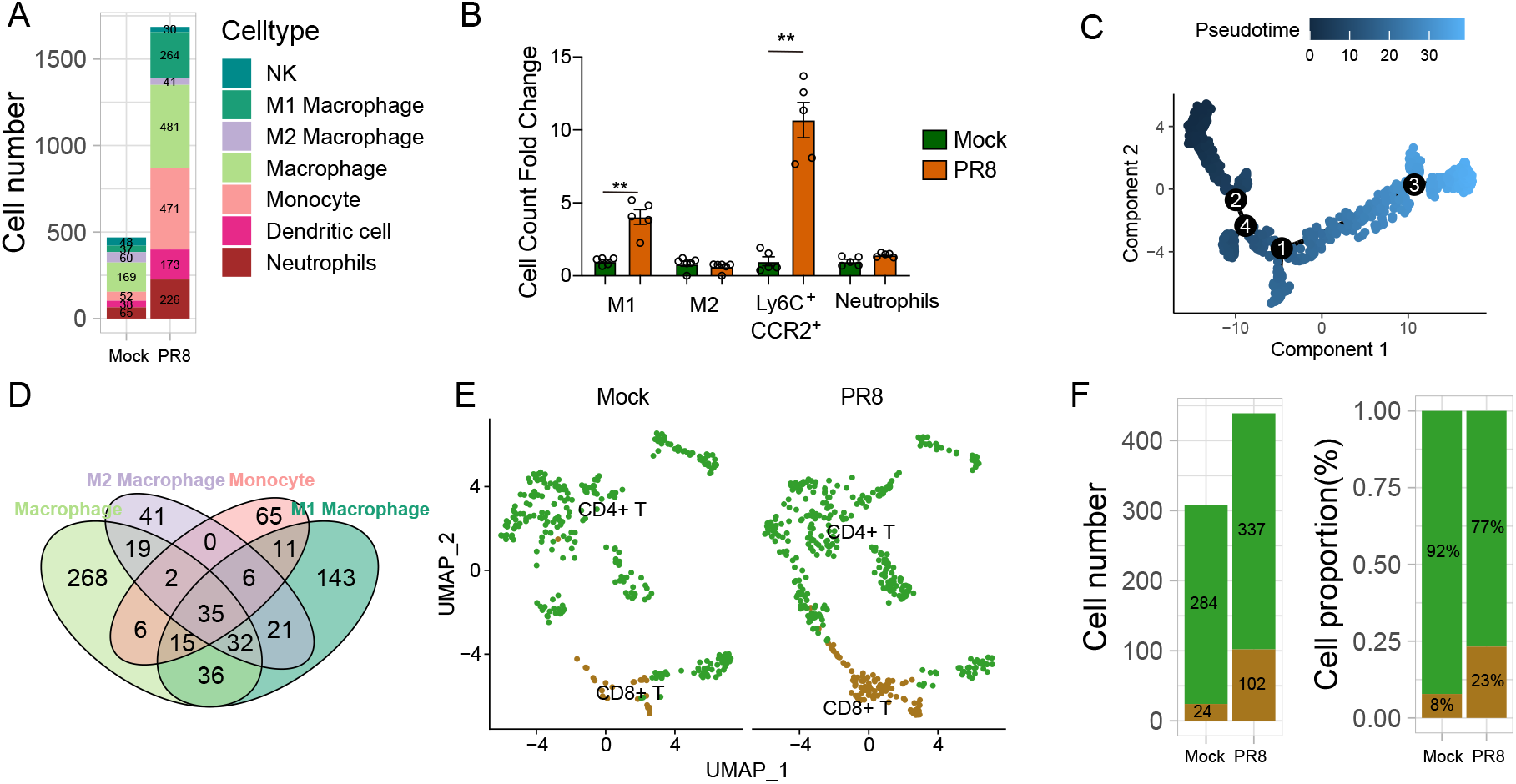
Cellular T cell landscapes in mock and IAV-infected lungs. Mice were infected with 250 PFU IAV. At 7 days post-infection, lungs were aseptically isolated and single-cell suspension was prepared for sc-RNAseq and flow cytometry. (A) Cell numbers of innate immune cells in the lungs of mock and IAV-infected mice. (B) FACS analysis of the number of lung cells from mock and IAV-infected mice at 7 days post-infection. Data shown as means ± SEM, reperesentative of 2 different experiments with n=5/group. Data analyzed by two-tailed Student’s *t*-test, ** P <0.01. (C) Single-cell trajectory through pseudotime constructed by reverse graph embedding of unbiased cluster analysis and dimensional reduction of single-cell transcriptomes. (D) Venn diagram showing the shared and unique differentially expressed genes in macrophages, M1 & M2 macrophages and monocytes between mock and IAV-infected mice. (E) UMAP projection embedding of T cells (CD4^+^ T cell and CD8^+^ T cells), color-coded by cell subsets as indicated. (F) Cell numbers and relative proportion of T cells among mock and IAV-infected lungs.

**Supplemental Figure 3.**
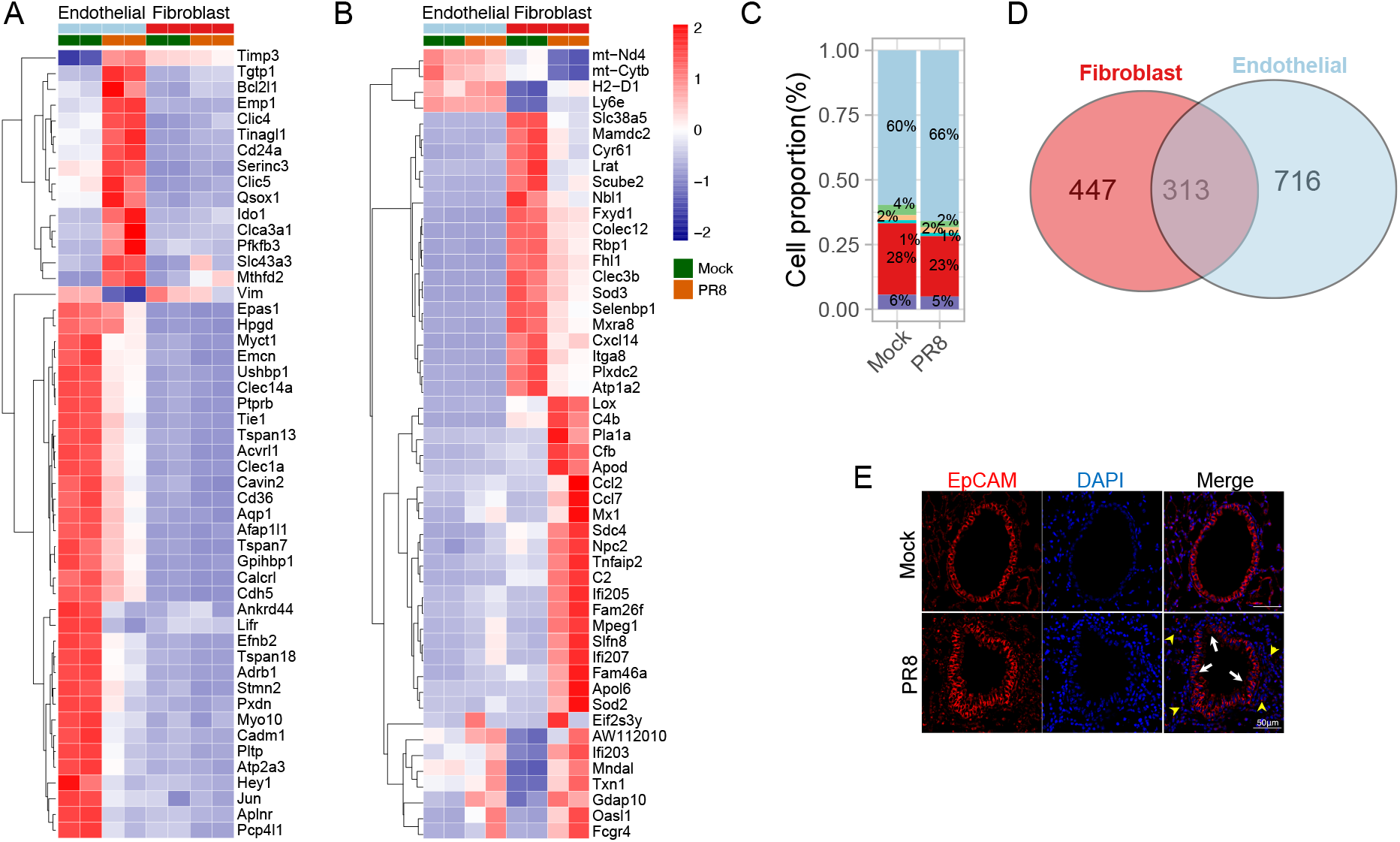
Non-hematopoietic cell landscapes in mock and IAV-infected lungs. Mice were infected with 250 PFU IAV. At 7 days post-infection, lungs were aseptically isolated and tissue was preserved for immunofluorescent staining or single-cell suspension was prepared for sc-RNAseq. (A, B) Heatmap depicting relative expression of top 50 unique differentially expressed (DE) genes in endothelial cells(A) and fibroblasts(B) between mock and IAV-infected lungs. (C) Cell relative proportion of barrier cells among mock and IAV-infected lungs. (D) Venn diagram shows the shared and unique differentially expressed (DE) genes in endothelial cells and fibroblasts between mock and IAV-infected lungs. (E) Immunofluorescent staining of EpCAM+ lung epithelial cells in mock and IAV-infected mice. IAV-infected mice show an infiltration of inflammatory cells in the peribronchial areas (yellow arrow heads), hyperplasia of airway bronchial epithelial cells associated with an extensive loss of the continuous integrity of epithelial layer (white arrows) compared to the airways of uninfected mice. EpCAM(red), nuclei were counterstained with DAPI (blue). Representative images of 2 different experiments with n=4. Images taken at X63 magnification, scale bar 50μm.

**Supplemental Figure 4.**
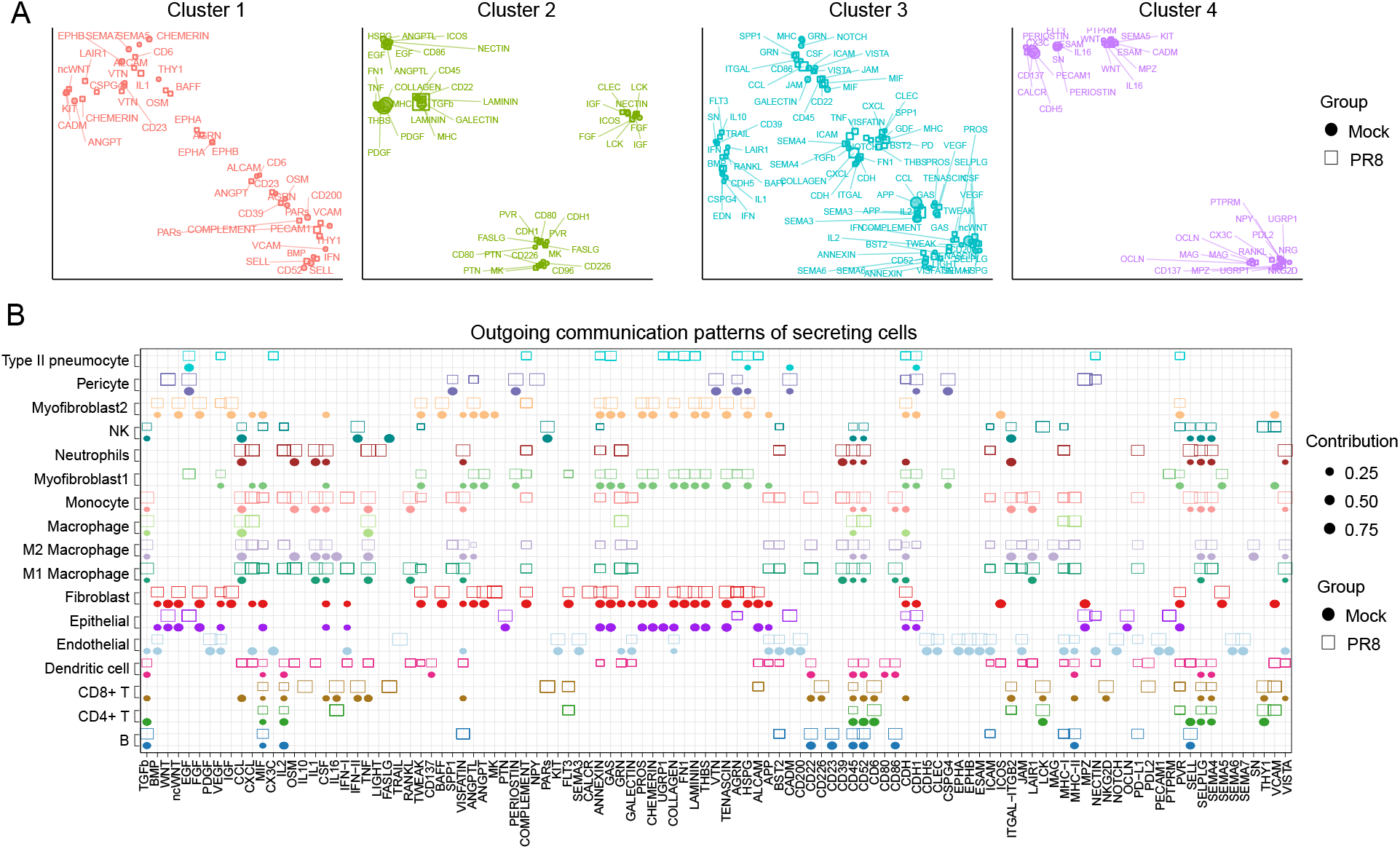
Cell-cell communications in mock and IAV-infected lungs. (A) Jointly projecting signaling pathways from mock and IAV infection onto shared two-dimensional manifold according to their structural similarity. Each dot represents the communication network of one signaling pathway. Dot size is proportion to the communication probability. Different colors represent different groups of signaling pathways. (B) The dot plot showing the comparison of outgoing signaling patterns of secreting cells between mock and IAV infection. The dot size is proportional to the contribution score computed from pattern recognition analysis. Higher contribution score implies the signaling pathway is more enriched in the corresponding cell group.

**Supplemental Figure 5.**
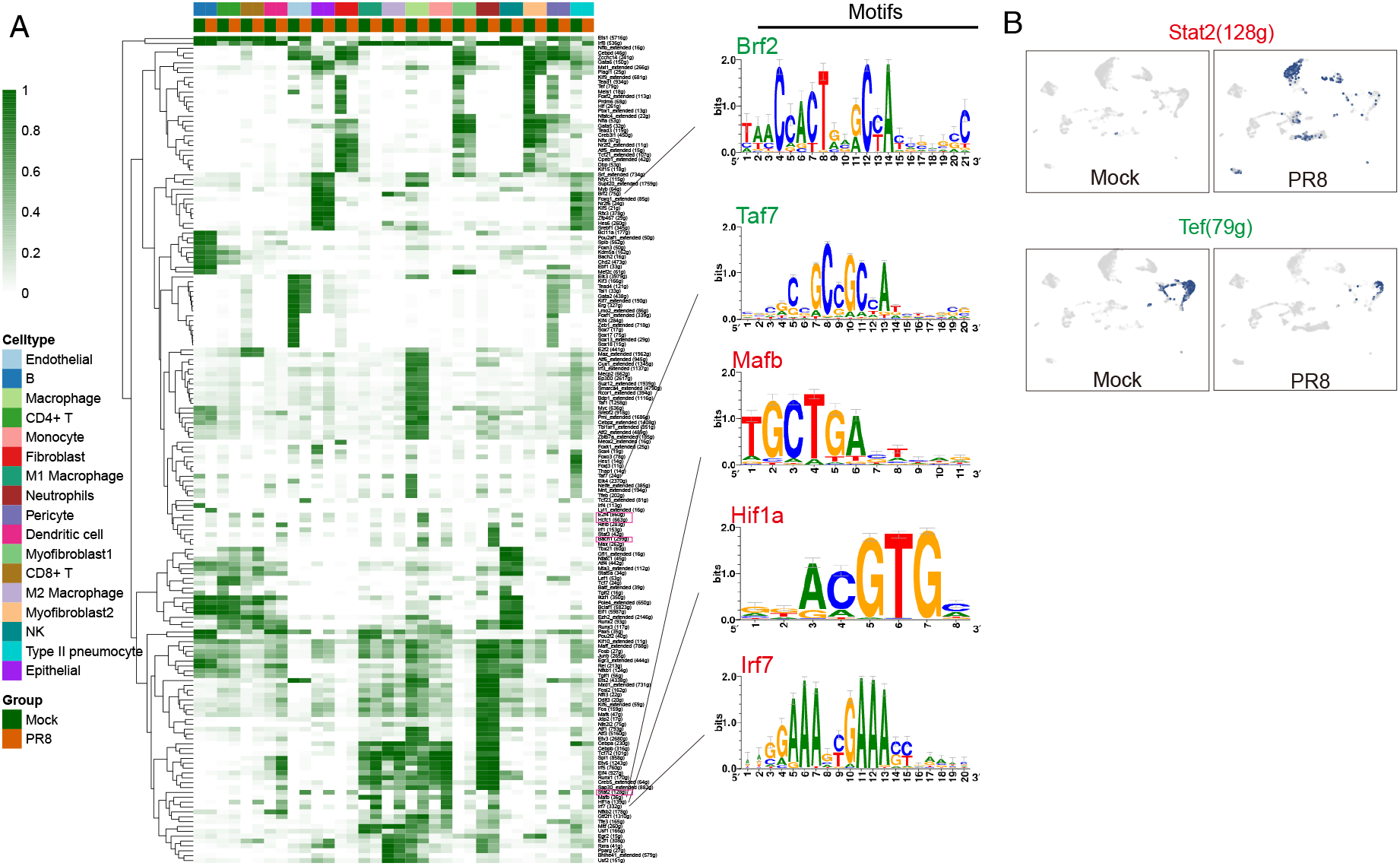
Transcriptional regulon in mock and IAV-infected lungs. (A) Heatmap of regulon activity analyzed by SCENIC with default thresholds for binarization. The enriched motifs from cisTarget for a selected set of regulons related to innate immune cells, generated within the SCENIC workflow. (B) UMAP projection of average binary regulon activity of *Stat2* and *Tef* in mock and IAV-infected lungs.

**Supplemental Figure 6.**
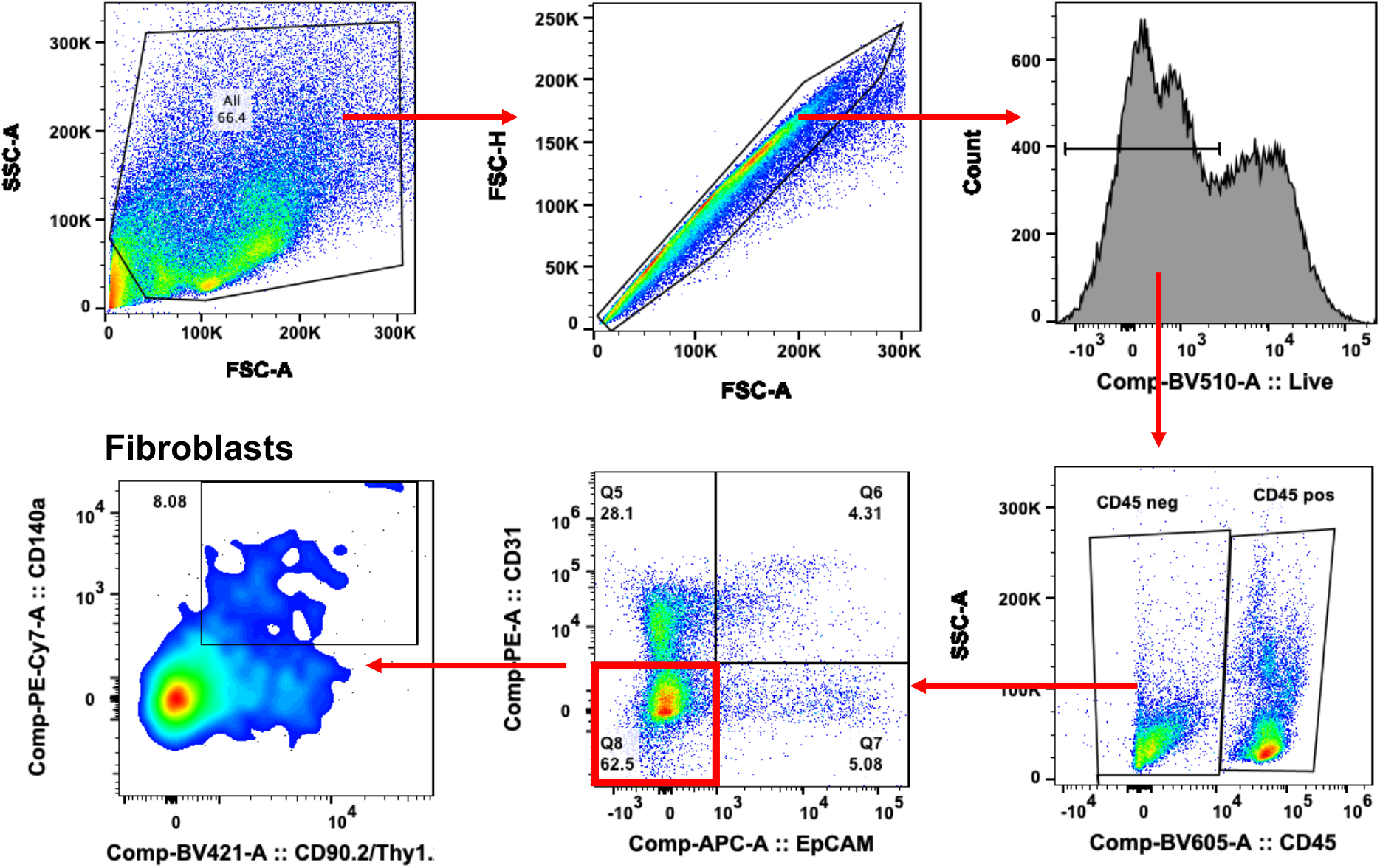
Flow cytometry gating strategy for lung fibroblasts. Mice were infected with 250 PFU IAV. At 7 days post-infection, lungs were aseptically isolated and single-cell suspension was prepared for flow cytometry. Lung fibroblasts were gated as CD45-CD31-EpCAM- CD90.2+ CD140a+ cells.

**Supplemental Figure 7.**
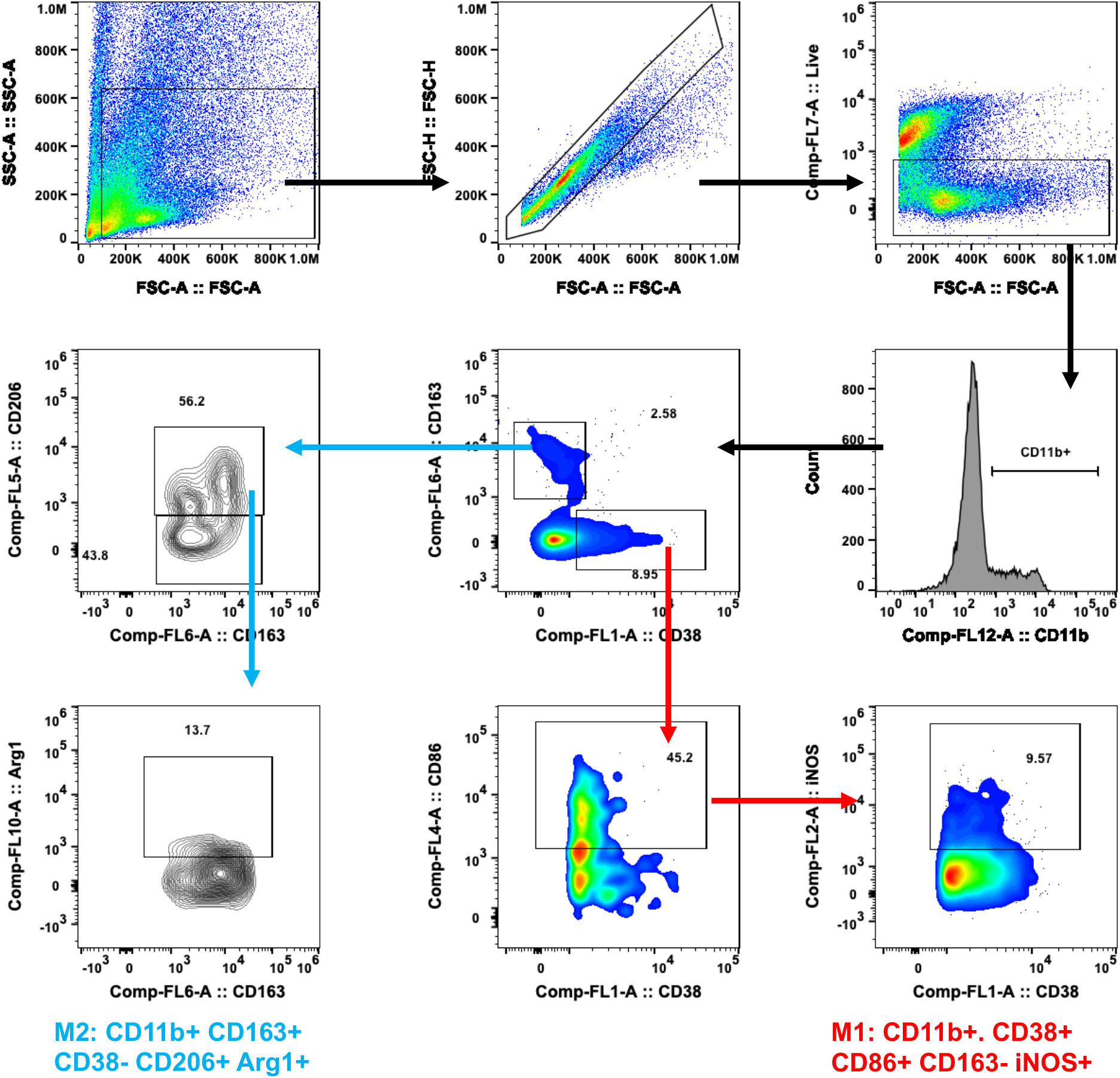
Flow cytometry gating strategy for M1 and M2 macrophages. Mice were infected with 250 PFU IAV. At 7 days post-infection, lungs were aseptically isolated and single-cell suspension was prepared flow cytometry. Cells were stained for fibroblasts, M1(CD11b+. CD38+ CD86+ CD163- iNOS+) macrophages and M2(CD11b+ CD163+ CD38-CD206+ Arg1+) macrophages.

## Notes

**Conflict of Interest.** All authors have read and approved the final version of the manuscript. The authors have declared that no conflict of interest exists.

### Competing Interest Statement

The authors have declared no competing interest.

